# Increased cGMP improves microvascular exercise training adaptations independent of endothelial nitric oxide synthase

**DOI:** 10.1101/2024.09.18.612717

**Authors:** Nathan C. Winn, David A. Cappel, Ethan D. Pollock, Louise Lantier, Jillian K. Riveros, Payton Debrow, Deanna P. Bracy, Joshua A. Beckman, David H. Wasserman

## Abstract

Impaired microvascular function is a hallmark of pre-diabetes. With development of atherosclerosis this impaired microvascular function can result in diminished capacity for ambulation and is a risk factor for Type 2 Diabetes. Dynamic changes in vascular tone are determined, in large part, by the eNOS/NO/cGMP axis. We used gain of function of the eNOS/NO/cGMP axis in diet-induced obese (DIO) mice and reduced function in lean mice to test the hypothesis that functionality of this vascular control mechanism parallels the benefits of an exercise training regimen. DIO mice have lower exercise capacity than lean mice and were used for pharmacological gain of function. The PDE-5a inhibitor – sildenafil – increases cGMP and was administered to DIO mice daily. In sedentary mice, we find that sildenafil does not improve exercise capacity. In contrast, it amplifies the microcirculatory effects of exercise training. Sildenafil synergizes with exercise training to improve performance during an incremental exercise test. Improved exercise performance was accompanied by increased skeletal muscle capillary flow velocity and capillary density measured via intravital microscopy. Loss of function was tested in lean mice hemizygous for endothelial cell (EC) specific eNOS creating an EC-eNOS knockdown (KD). EC-eNOS KD decreases capillary density and exercise tolerance in sedentary mice; however, it did not prevent exercise-training induced improvements in endurance capacity. These data show that 1) increasing cGMP with sildenafil enhances microcirculatory function and exercise work tolerance that results from training; 2) eNOS KD does not prevent the microcirculatory or improvements in exercise tolerance with training. PDE-5a inhibitors combined with physical exercise are a potential mechanism for improving ambulation in patients with circulatory limitations.

## INTRODUCTION

Capillaries of the microcirculation are the site of molecule exchange between blood and the cells of perfused tissues (1, 2). Sustained increase in metabolic rate requires vascular delivery to maintain high metabolic activity as tissue stores of oxygen are limited. Peripheral artery disease (PAD) and diabetes cause microcirculatory defects that limit the exchange of bloodborne proteins and metabolites and impede limb muscle capillary blood flow. This can lead to significant limb ischemia that results in reduced ambulation, intermittent claudication, and a reduced quality of life. Endothelial- and non-endothelial derived factors determine vascular conductance at arterioles. Endothelial nitric oxide synthase (eNOS) catalyzes the production of nitric oxide (NO) from L-arginine. NO activates guanylate cyclase in the surrounding smooth muscle to produce cGMP which stimulates protein kinase G (PKG). The result of cGMP-PKG signaling is smooth muscle relaxation, vasodilation, and angiogenesis (3, 4). The magnitude and duration of cGMP concentrations is reduced by hydrolysis of cGMP by a cGMP-dependent phosphodiesterase. Phosphodiesterase-5a (PDE-5a) is the predominant phosphodiesterase in vascular smooth muscle. Sildenafil is a highly selective inhibitor of PDE-5a that increases cGMP by inhibiting cGMP hydrolysis. This results in a decrease in vascular resistance. PDE-5a inhibitors are approved for the treatment of erectile dysfunction; however, it may also have therapeutic benefits in treatment of primary pulmonary hypertension (5), as well as limiting deficits in learning and memory processes associated with neurodegeneration of aging (6, 7).

Studies show that chronic PDE-5a inhibition enhances insulin sensitivity in rodents (1, 8) and humans (9). Capillary density is closely associated with skeletal muscle insulin sensitivity (5,6). Contracting skeletal muscle requires accelerated microcirculatory nutrient delivery and removal of metabolic byproducts. Dynamic changes in vascular tone are determined, in large part, by the eNOS/NO/cGMP axis which during exercise is accelerated causing smooth muscle relaxation of skeletal muscle arterioles (10) and accelerated blood flow to working muscle. From a clinical perspective, reduction in walking capacity is a common adverse sequela of PAD. Exercise rehabilitation improves ambulation in these patients (11). However, ischemic pain associated with ambulatory activity can limit its utility. Amplification of the eNOS/NO/cGMP axis may cause physical activity-induced improvements in microvascular and physical function, while impaired activity of this axis has been shown to adversely affect a single bout of exercise.

This study used gain and partial loss of eNOS/NO/cGMP function approaches to address how microvascular function affects the response to regular physical activity. The gain of function approach used pharmacological administration of PDE5a inhibitor, sildenafil, in diet-induced obese (DIO) mice that are known to have impaired endothelium-dependent vasodilation (12-14). The partial loss of function approach utilized mice with an inducible endothelial cell (EC) specific eNOS hemizygous deletion (eNOS+/-) to test whether effects of physical activity are attenuated by a decrease in eNOS/NO/cGMP axis in otherwise lean healthy mice. Results show that gain of function studies using pharmacological inhibition of PDE-5a improves microvascular function and augments the effects of regular physical activity in DIO mice. In contrast, partial loss of function studies using EC-specific knockdown (KD) of eNOS+/-does not impair the effects of physical activity.

## METHODS

Procedures involving mice were approved in advance and carried out in compliance with the Vanderbilt University Institutional Animal Care and Use Committee. Vanderbilt University is accredited by the Association for Assessment and Accreditation of Laboratory Animal Care International. The environmental temperature at which mice were housed and experiments were performed was between 21-23°C.

### Animals

All mice were on the C57BL/6J strain and fed standard chow diet (5001 Laboratory Rodent; LabDiet) or 60% fat diet (HFD, Research Diets, #D12492, 5.21 kcal/g food). *Protocol 1*: Mice hemizygous for the lox-p flanked gene *Nos3* (eNOS) were crossed with mice expressing the stem cell leukemia (SCL) promoter driven tamoxifen-inducible Cre recombinase-estrogen receptor fusion protein (CreER) transgene (15, 16). This construct causes EC-specific gene deletion (17, 18). Mice heterozygous for the lox-p flanked gene Nos3 without the CreER transgene were used as controls. Both mouse lines were administered tamoxifen (TMX; Sigma-T5648) reconstituted in corn oil vehicle (Sigma-C8267) at 2 mg/kg body weight for 5 days to generate an EC-specific eNOS knockdown or WT littermate controls. EC eNOS+/- and WT mice underwent 5 weeks of exercise training or remained sedentary. *Protocol 2*: C57BL/6J were fed HFD for 8 weeks to evoke DIO. Sildenafil (6 *µ*g•g^-1^ body weight; PHR1807, Millipore Sigma) versus vehicle (volume matched injection) was delivered via subcutaneous injection two times per day (∼10 hours between injections) for 5 weeks. Vehicle injections consisted of 0.9% saline.Chow fed age-matched mice were used as lean controls. After 8 weeks on diet, mice either remained sedentary or underwent exercise training for 5 weeks (detailed below), while being treated with Sildenafil or vehicle. Chow fed control mice were sedentary or underwent exercise training, but were not treated with Sildenafil. Another cohort of mice were fed HFD for 8 weeks to induce DIO and then were treated twice daily with vehicle injections for 5 weeks. After 5 weeks, mice were acutely injected with Sildenafil (6 *µ*g•g^-1^ body weight) or vehicle (volume matched 0.9% saline) to test whether acute treatment with sildenafil improves hemodynamics and exercise tolerance.

### Incremental Exercise Stress Test and Exercise Training

The stress tests were conducted on a single-lane treadmill from Columbus Instruments beginning at a speed of 10 m·min^-1^. Speed was increased by 4 m·min^-1^ every three minutes until exhaustion. Mice exercised at 40% of their maximum speed (measured during the stress test) for 30 min, 5 days per week for 5 weeks. This results in an exercise training volume of 150 minutes of moderate intensity exercise per week.

### Intravital Microscopy

Intravital microscopy was performed in mice as previously described (19-21). Studies were performed under isoflurane anesthesia (SomnoSuite; Kent Scientific). Doses of 2 and 1.5% isoflurane were used for induction and maintenance of anesthesia, respectively. The lateral gastrocnemius was exposed for visualization by trimming away skin and fascia and then placed on a glass coverslip immersed in 0.9% saline. Body temperature of 37°C was maintained using a homeothermic heating blanket and temperature probe (Harvard Apparatus). Approximately 8 mg/kg (100 μL injection volume) of a tetramethylrhodamine (TMR)-labeled 2-megadalton (mDa) dextran (Thermo Fisher) was infused through the jugular vein catheter. A 2 mDa molecule remains in the lumen of capillaries with high resistance endothelial walls and used as to identify microcirculatory structure. Imaging began at t=0 min and continued until 10 minutes after the TMR bolus and insulin bolus, respectively. The focal plane for imaging was maintained using the Perfect Focus System on the Nikon Eclipse Ti-E (Nikon Instruments). Plasma fluorescence was excited using a Sola Light Engine LED lamp (Lumencore) and visualized with a Plan Apo 10× objective (Nikon Instruments). Images were recorded at 100 fps using a 10 ms exposure time with a Flash 4.0 sCMOS camera (Hamamatsu). High frame rate imaging was achieved by directly streaming time-lapse experiments to computer RAM using a Camlink interface (Olympus). Videos underwent real-time 4 × 4 pixel binning during acquisition to improve signal/noise ratio, and image brightness was manually adjusted post hoc in NIS Elements (Nikon Instruments) to ensure an average pixel intensity of approximately 0.5 with a minimal extent of over/underexposed regions prior to export in a MATLAB-readable format. Capillary flow velocity, hematocrit, and density were quantified as previously described (22). Briefly, 5 s videos of the gastrocnemius microcirculation acquired at 100 frames·s^-1^ were processed to remove motion artifacts, identify in-focus capillaries, and track the motion of red blood cells (RBCs) based on the shadows they produce in plasma fluorescence. Perfusion metrics included mean capillary flow velocity (MFV; µm·s^-1^), and perfusion heterogeneity index (PHI; a unitless measure of spatial flow variability). PHI was defined as the natural log of *V*□_max_ - *V*□_min_ divided by the MFV. *V*□_max_ and *V*□_min_ represent velocity in the fastest and slowest flowing capillaries, respectively. Five fields of view were captured for each mouse, and MFV and PHI represent an average of all five acquisitions.

### cGMP analysis

Circulating cGMP levels were determined from arterial plasma samples using a cGMP immunoassay kit (#CG200, Millipore Sigma). Arterial blood was collected 5 hours from the last dose of sildenafil.

### Immunoblotting

Immunoblotting was performed on gastrocnemius muscle in EC eNOS+/- and WT mice. Skeletal muscle was excised. Muscle tissue was minced and digested in HBSS containing 5% fetal bovine serum, 2U/mL Dispase, and 2mg/mL collagenase type II for 35 minutes at 37°C. Digested muscle tissue was then vortexed, filtered through 100 μm filters and centrifuged at 400xg to pellet cells. Supernatant were discarded and cell pellets were incubated with ACK buffer to lyse red blood cells. Cells were filtered through 35 μm filters and incubated with CD31 magnetic conjugated bead antibodies (#130-097-418, Miltenyi Biotec). Positive selection for CD31+ cells was performed using magnetic separator columns (#130-042-201, Miltenyi Biotec). CD31- and CD31+ cells were flash frozen and stored at -80°C. Triton X-100 CD31+ and CD31-cell lysates (5-15 µg/lane) were analyzed by western blot for proteins and eNOS (#32027; 1:500; Cell Signaling) and GAPDH (#97166; 1:1000; Cell Signaling). After incubation with appropriate HRP-conjugated secondary antibodies (rabbit anti-IgG, #7074, mouse anti-IgG #7076, Cell Signaling) bands were detected via chemiluminescence. Intensity of individual bands were quantified using Image Lab^TM^ (version 6.0.0, Bio-Rad Laboratories, Inc.), and expressed as a ratio to GAPDH.

### Statistical Analysis

Student’s t-tests were run for between group comparisons. If data did not follow a Gaussian distribution, non-parametric Mann-Whitney tests were used to determine

### statistical significance

In experiments that contained more than two groups, one-way analysis of variance (ANOVA) or two-way ANOVA models were used with pairwise comparisons using Tukey or Sidak correction. Brown-Forsythe correction was applied to groups with unequal variance. Data are presented as mean ± standard error (SE). A p value of <0.05 was used to determine significance.

## RESULTS

### EC eNOS knockdown does not prevent improvements in microcirculation or exercise tolerance with exercise training

We tested whether a decrease in eNOS/NO/cGMP axis in otherwise lean healthy mice blunts microvascular function and exercise tolerance with training. Mice with an inducible KD of EC eNOS were used for this purpose. To KD EC eNOS, mice expressing the EC specific SCL promoter Cre ERT transgene or controls not expressing SCL cre were administered tamoxifen for 5 consecutive days. Five weeks later, mice were euthanized to quantify the magnitude of eNOS KD in EC enriched skeletal muscle. Since skeletal muscle is comprised of heterogenous cell types, we used magnetic conjugated bead selection to enrich for ECs (defined by CD31+ expression) (**Fig. 2A**). eNOS expression was restricted to CD31 expressing cells with no eNOS expression detected in the CD31 negative fraction (**Fig. 2A**). This confirms the specificity of EC isolation. Using the bead-based EC enrichment strategy, we found that mice hemizygous for EC-specific eNOS have a ∼50% eNOS knockdown (**Fig. 2C&D**). WT and EC eNOS+/-mice had similar body weight, % fat mass, and % lean mass (**Fig. 2E&F**). Together, these data confirm quantitative KD of eNOS in CD31+ ECs.

**Figure 1.**
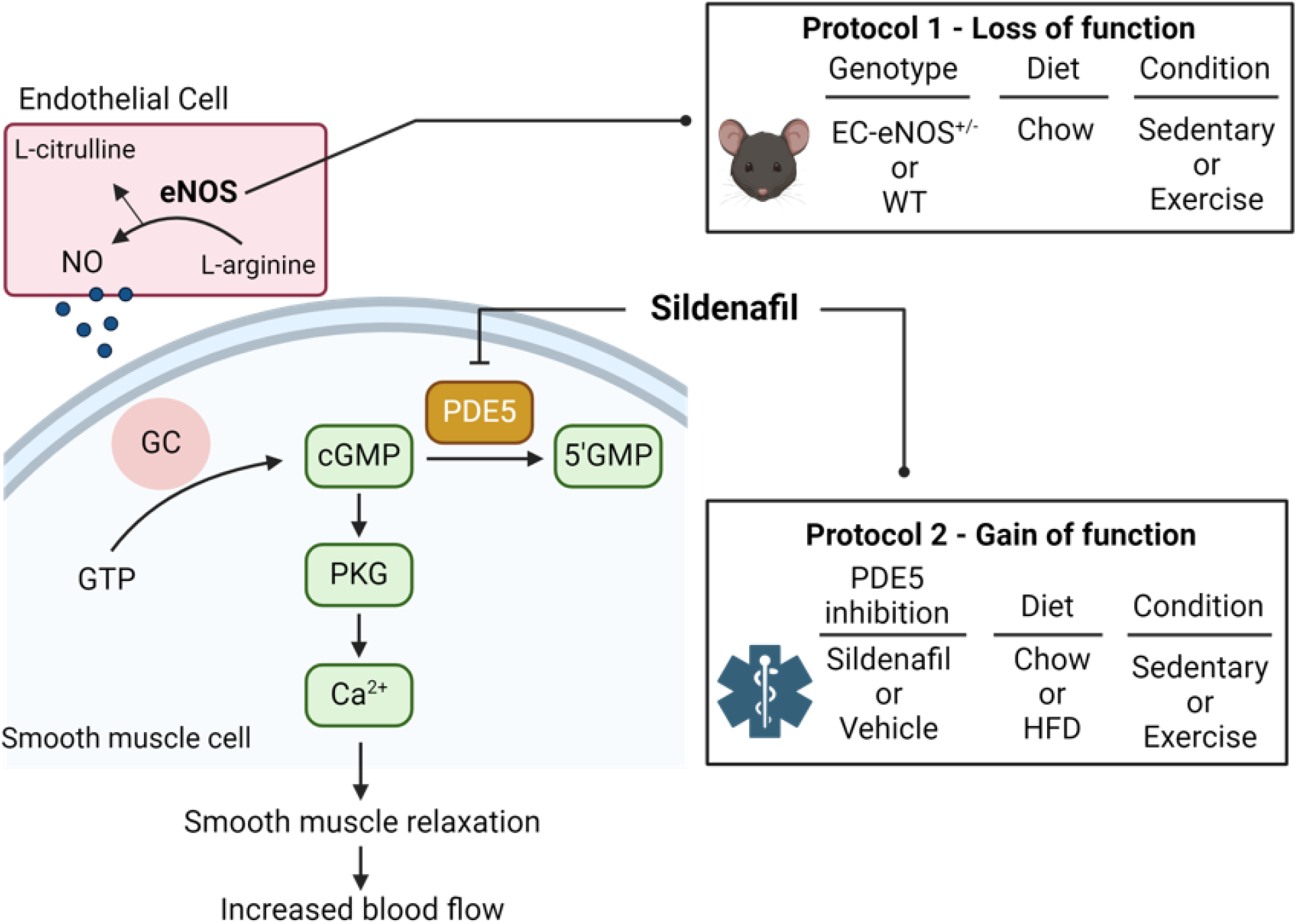
Summary of the eNOS/cGMP/PKG pathway and experimental cohorts. The activation of eNOS leads to increased NO production from endothelial cells (EC). NO activates guanylate cyclase in the surrounding smooth muscle to produce cGMP which stimulates protein kinase G (PKG). The result of cGMP-PKG signaling is smooth muscle relaxation, vasodilation, and angiogenesis. The magnitude and duration of cGMP concentrations is reduced by hydrolysis of cGMP by a cGMP-dependent phosphodiesterase. Phosphodiesterase-5a (PDE-5a) is the predominant phosphodiesterase in vascular smooth muscle. Sildenafil is a highly selective inhibitor of PDE-5a that increases cGMP by inhibiting cGMP hydrolysis. In protocol 1 a loss of function mutant mouse with EC-specific loss of an eNOS allele is used to test the hypothesis that a reduction in eNOS blunts exercise training induced adaptations in the microvasculature. In protocol 2, C57BL/6J were fed HFD for 8 weeks to evoke diet-induced obesity (DIO). Sildenafil (6µg•g-1 body weight) versus vehicle (volume matched injection) was delivered via subcutaneous injection two times per day for 5 weeks with or without exercise training.

**Figure 2.**
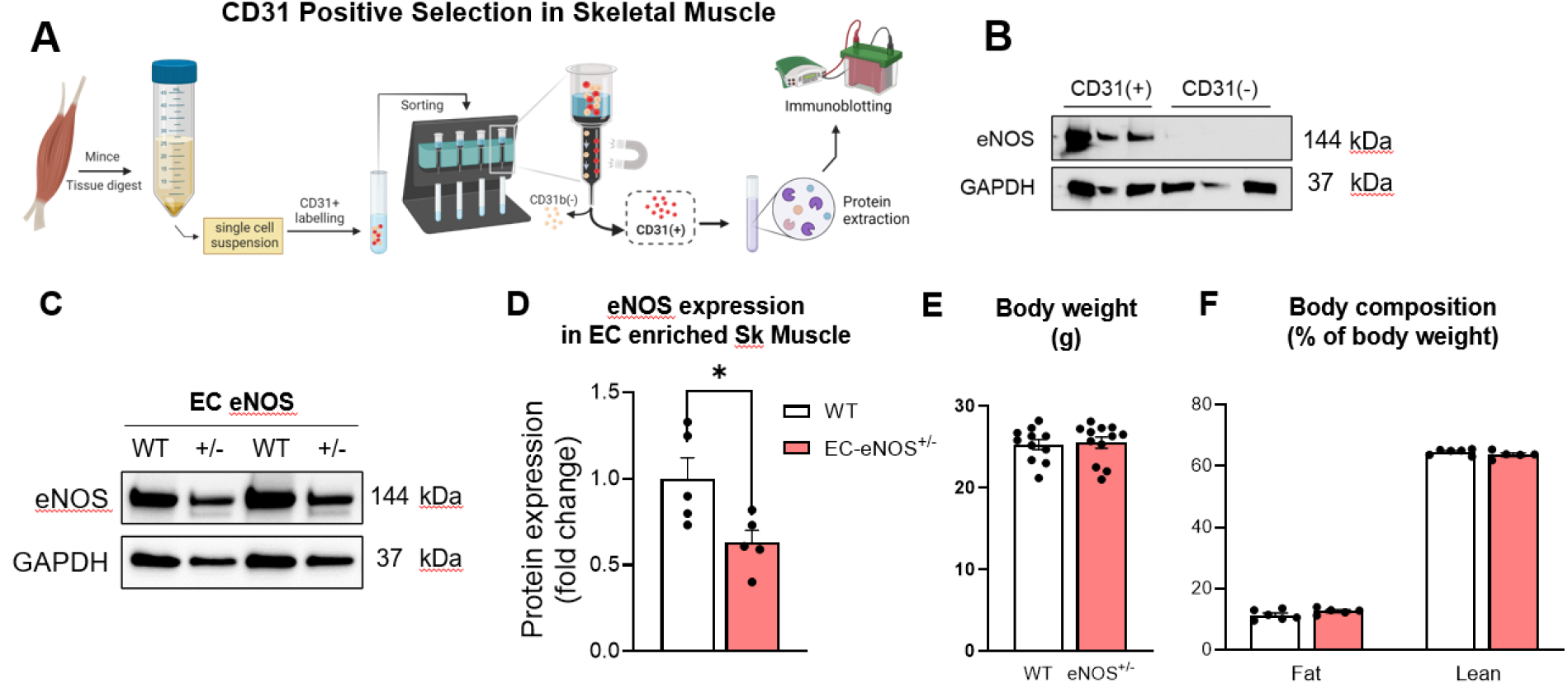
Confirmation of EC eNOS knockdown and body composition. A) Gastrocnemius muscle was digested and single cell suspension generated. CD31 conjugated beads were used to enrich for endothelial cells. The CD31+ and CD31-fractions were prepped for protein extraction and B) immunoblotting was performed for eNOS and GAPDH. These data show that only the CD31+ fraction expresses eNOS protein. C) CD31 enrich muscle ECs isolated from WT and EC-eNOS+/-mice were immunoblotted for eNOS and GAPDH. D) Densitometry of eNOS immunoblot relative to WT control mice. E) Body weight and F) body composition analysis in WT versus EC-eNOS+/-mice. Sk muscle, skeletal muscle. EC, endothelial cell. Two tail student t test were run to compare groups. n=5-12 mice per group. Data are mean ± SE. *p<0.05.

WT and EC eNOS+/-mice were randomized to exercise training or remained sedentary for 5 weeks (**Fig. 3A**). Exercise tolerance in response to an exercise stress test was lower in sedentary eNOS+/-compared to WT sedentary mice (**Fig. 3B**). Exercise training led to similar increases in exercise tolerance in WT and EC eNOS+/-mice. Intravital microscopy was performed approximately 36 hours after the last exercise bout in gastrocnemius muscle. No differences in skeletal muscle capillary blood flow velocity existed between groups **(Fig. 3C**). The capillary density within the FOV was reduced in sedentary eNOS+/-compared to WT mice (**Fig. 3D**). Although there were no differences in capillary blood flow, the flow heterogeneity was decreased with exercise training in both WT and eNOS+/-mice (**Fig. 3E**). Hematocrit variability reflects the movement of particulates in a capillary bed, which was similarly reduced between genotypes with exercise training (**Fig. 3F**). Collectively, these data show that partial loss of EC eNOS does not prevent improvements in microcirculatory function or exercise tolerance with training in lean mice.

**Figure 3.**
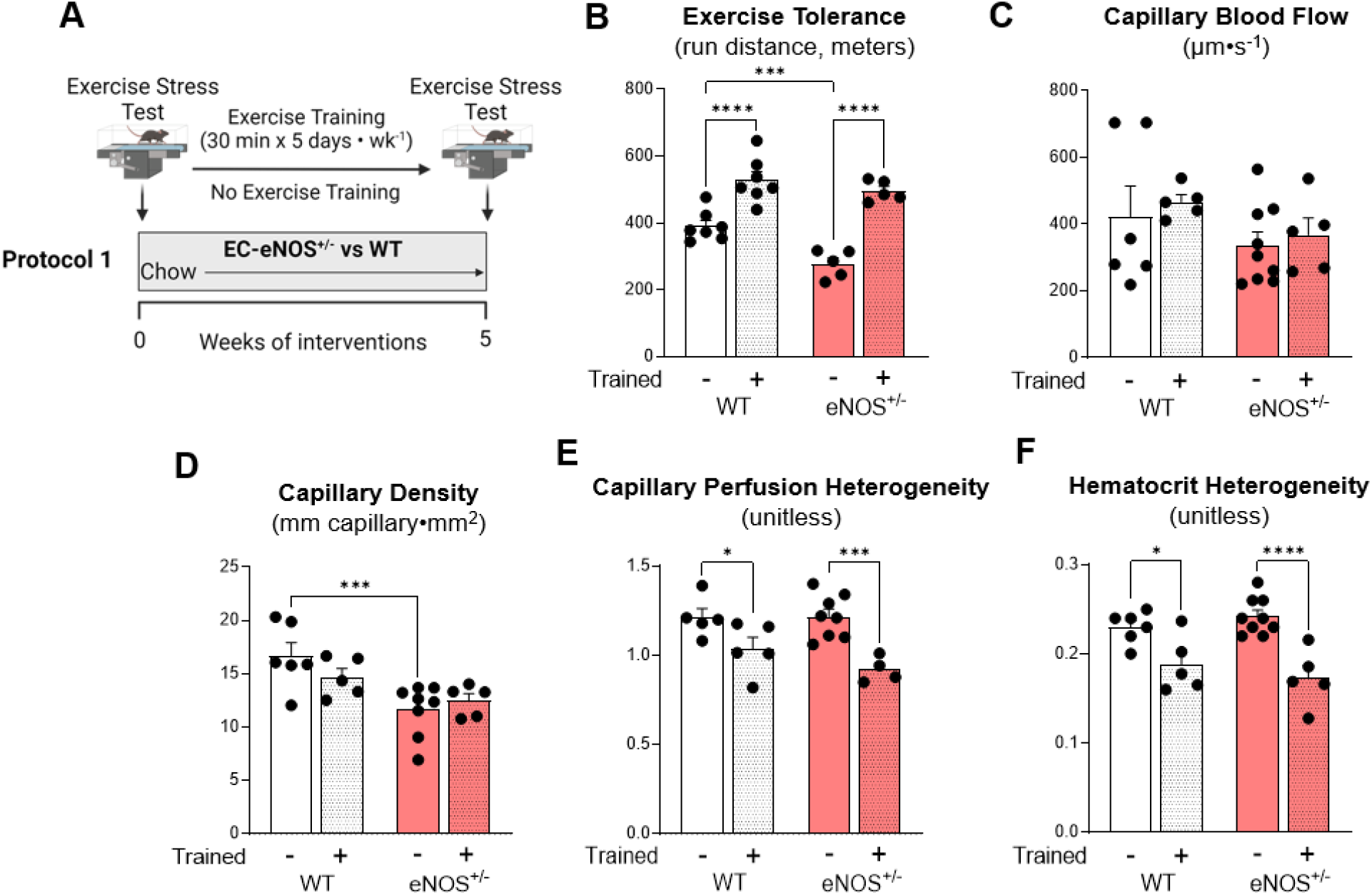
EC eNOS knockdown decreases capillary density and exercise tolerance in sedentary mice, but does not prevent training-induced adaptations in microvascular hemodynamics. **A**) EC eNOS+/- and WT mice underwent a baseline exercise stress test to determine exercise training intensity. Mice exercised 30 minutes per day, 5 days per week for 5 weeks on a treadmill. **B**) Exercise stress test after the 5 week training period. Mice ran on a treadmill with increasing intensity every 3 minutes until the animals could no longer maintain the running speed. The greater the distance travelled the greater the exercise tolerance. Post exercise training microvascular hemodynamics were measured using intravital microscopy on the gastrocnemius muscle to determine **C**) capillary blood flow rate, **D**) capillary density, and **E**) perfusion heterogeneity, and **F**) hematocrit heterogeneity. n=4-9 mice per group. Student t tests were run for two group comparisons. Two-way ANOVA with genotype and training as factors were run for panels B-F Significance was accepted when p<0.05. *p<0.05, **p<0.01, ***p<0.001, ****p<0.0001

### Acute PDE-5 inhibition increases capillary blood flow but does not improve exercise tolerance in DIO mice

DIO is associated with microvascular dysfunction (23, 24). We tested the hypothesis that acute administration of sildenafil enhances microvascular function and exercise performance in DIO mice. After 8 weeks of DIO, mice received two subcutaneous injections of sildenafil 16 hours and 5 hours prior to an exercise stress test or measurements of microvascular hemodynamics (**Fig. 4A**). Despite a ∼5-fold increase in arterial cGMP (**Fig. 4B**) acute treatment with sildenafil did not improve exercise tolerance (**Fig. 4C**). Capillary blood flow was increased by ∼30% in acute sildenafil treated mice **(Fig. 4D)**. However, no change in perfusion heterogeneity or hematocrit heterogeneity were detected **(Fig. 4E-F)**. These data suggest that acute sildenafil enhances capillary flow in muscle but this does not augment exercise performance in DIO mice.

**Figure 4.**
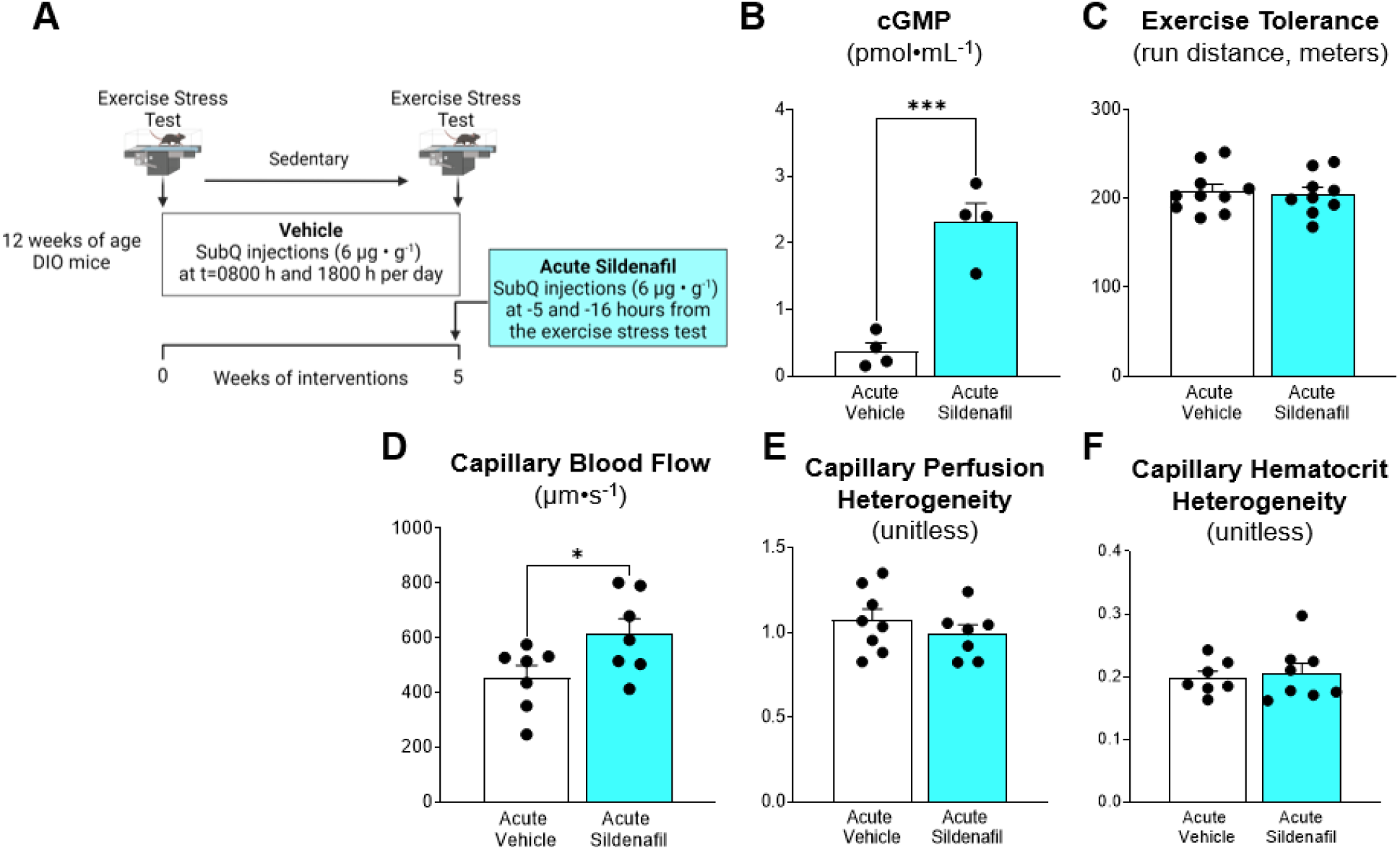
Acute administration of sildenafil enhances microvascular blood flow, but does not augment exercise tolerance in diet-induced obese mice. **A**) C57BL/6J were fed HFD for 8 weeks to evoke DIO. Mice underwent exercise training or no training for 5 weeks while receiving vehicle treatment. After 5 weeks of training, Sildenafil (6 µg•g^-1^ body weight) or vehicle was administered -16 hours and -5 hours before an exercise tolerance test. **B**) Arterial cGMP levels. **C**) Exercise tolerance was performed with total distance used as a readout of peak exercise capacity. Intravital microscopy was performed on gastrocnemius muscle as detailed in the Methods to assess microvascular hemodynamics. **D**) Capillary blood flow, **E**) capillary perfusion heterogeneity and **F**) capillary hematocrit heterogeneity was determined following acute sildenafil or vehicle administration. Two tail student t test were run to compare groups. n=4-10 mice per group. Data are mean ± SE. *p<0.05; ***p<0.001.

### Chronic PDE-5 inhibition amplifies exercise training-induced improvements in microvascular function and exercise tolerance

Exercise training causes adaptive responses to the microcirculation that typically result in gain of function in the eNOS/NO/cGMP axis. These adaptations to training may be blunted in DIO. We hypothesized that chronic treatment with sildenafil would synergize with exercise training to enhance microcirculatory hemodynamics and exercise tolerance. Lean and DIO mice were randomized to 5 weeks of treadmill exercise training or placed on the stationary treadmill belt and remained sedentary (**Fig. 5A**). The DIO groups received vehicle or sildenafil twice per day over the 5 week period. Prescreen exercise stress test results were used to ensure mice in sedentary and trained mice had initial equivalent exercise capacity and to determine exercise intensity in trained groups. As expected, DIO mice had reduced exercise tolerance compared with chow controls (**Fig. 5B**). After the 5 weeks of intervention arterial cGMP levels were increased ∼10-fold in mice chronically treated with sildenafil (**Fig. 5C**). DIO mice had greater body mass and fat mass than lean groups regardless of sildenafil treatment (**Fig. 5D**). Exercise training decreased body weight gain in DIO mice. Body fat percentage was lower in trained mice treated with sildenafil (**Fig. 5F**). The percent lean mass was lower in DIO mice than lean mice.Exercise training preserved lean mass in vehicle (p=0.056) and sildenafil treated groups (**Fig. 5E**).

**Figure 5.**
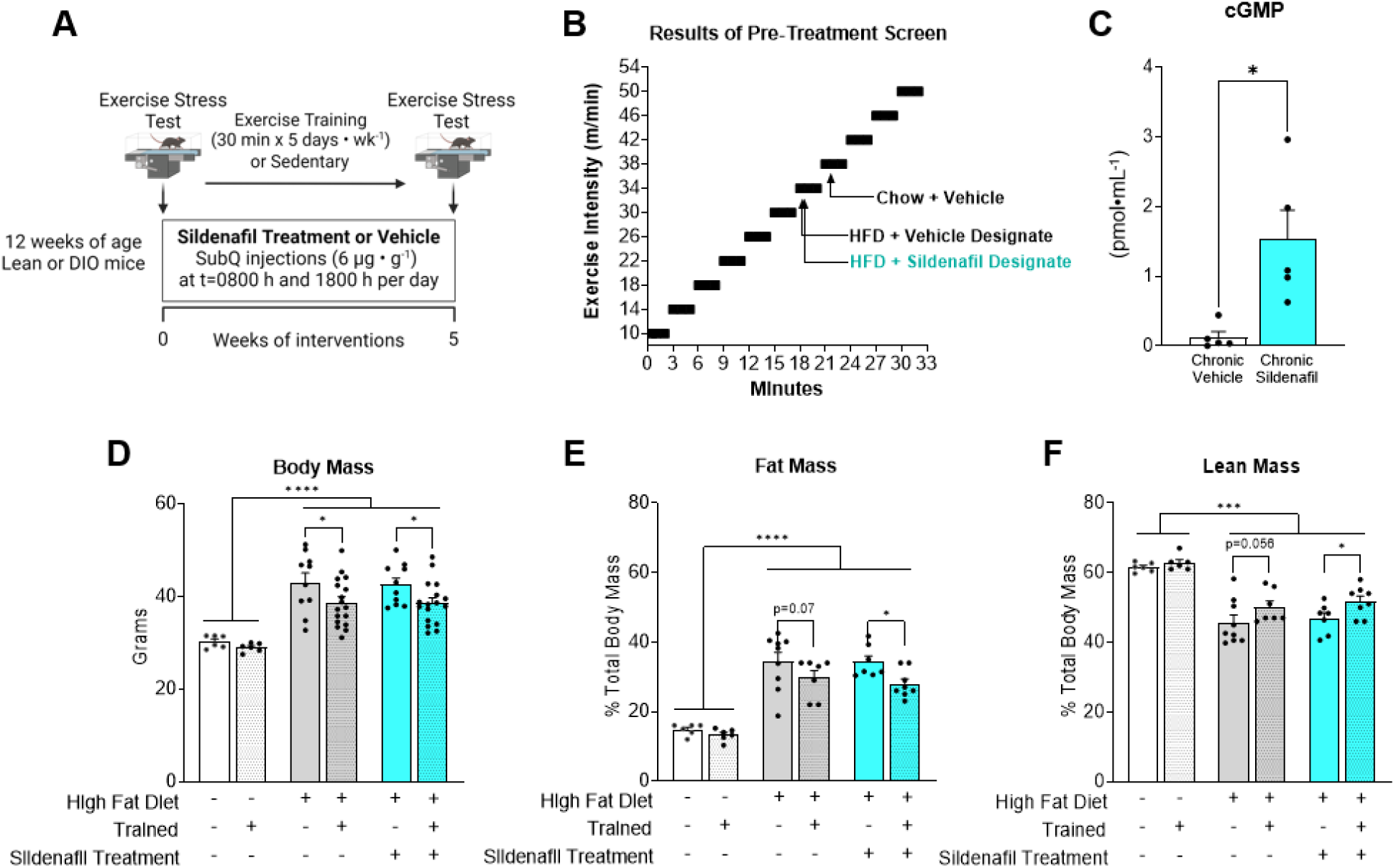
Exercise training protocol and body composition in trained versus sedentary mice receiving chronic vehicle or sildenafil treatment. **A**) C57BL/6J were fed HFD for 8 weeks to evoke DIO. Sildenafil (6 µg•g^-1^ body weight) versus vehicle (volume matched injection) was delivered via subcutaneous injection two times per day for 5 weeks while mice underwent exercise training or remained sedentary. **B**) Exercise tolerance test were performed after 8 weeks of DIO to determine the exercise intensity of the training period. **C**) Arterial cGMP levels in vehicle versus sildenafil groups. **D**) Body mass, **E**) body fat percentage and **F**) percent lean mass was determined after the 5 week training/sedentary period. Two way ANOVA with group (Lean vehicle, DIO vehicle, DIO Sildenafil) and training (Sedentary versus Exercise) as factors were run with Tukey correction for multiple comparisons. Data are expressed as mean ± SE n=5-16 mice per group. *p<0.05; ***p<0.001; ****p<0.0001.

Consistent with prescreen results, lean mice had greater exercise tolerance than DIO mice when compared within sedentary or trained conditions (**Fig. 6A**). Exercise training improved exercise tolerance in all groups. Chronic sildenafil treatment enhanced the improvement in exercise capacity with exercise training (**Fig. 6A**). Capillary blood flow velocity did not increase with exercise training in lean or vehicle DIO mice. However, sildenafil caused a 40% increase in capillary blood flow in trained DIO mice (**Fig. 6B**). Exercise trained mice had an increase in capillary density per FOV (exercise effect, p=0.004), with no differences between vehicle or sildenafil (**Fig. 6C**). Chronic sildenafil decreased the perfusion heterogeneity and the hematocrit heterogeneity (**Fig. 6D&E**). These data show that chronic treatment with sildenafil amplifies exercise training induced improvements in exercise tolerance and microcirculatory hemodynamics.

**Figure 6.**
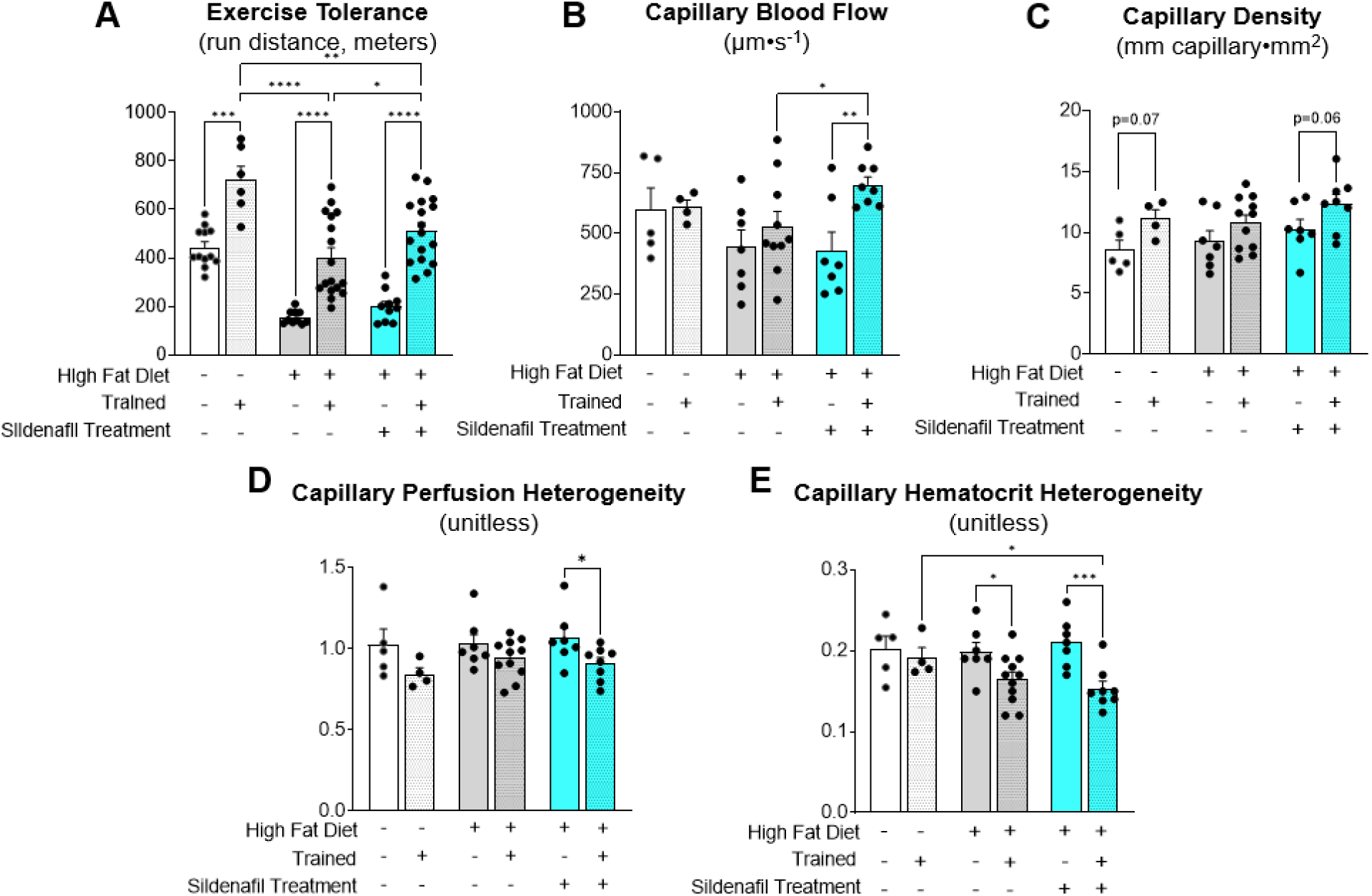
Chronic PDE-5 inhibition amplifies exercise training-induced improvements in microvascular function and exercise tolerance. A) Exercise tolerance was determined in lean and DIO mice with or without exercise training. DIO mice were treated with vehicle or sildenafil during the 5 week training or sedentary period. Exercise tolerance was determined as the peak run distance during an incremental exercise test. Intravital microscopy was used to determine microvascular hemodynamics in gastrocnemius muscle 36-48 hours after the last exercise bout. Microvascular readouts included B) capillary blood flow, C) capillary density, D) capillary perfusion heterogeneity, and E) capillary hematocrit heterogeneity. Two way ANOVA with group (Lean vehicle, DIO vehicle, DIO Sildenafil) and training (Sedentary versus Exercise) as factors were run with Tukey correction for multiple comparisons. n=4-17 mice per group. Data are expressed as mean ± SE. *p<0.05; **p<0.01; ***p<0.001; ****p<0.0001.

## DISCUSSION

This study used pharmacological gain of function and genetic loss of function approaches to test the hypothesis that the eNOS/NO/cGMP axis is central to exercise training adaptations in microcirculatory function and exercise capacity. We show that a single eNOS allele in ECs is sufficient for the microcirculatory adaptations and improvements in exercise tolerance with training in otherwise healthy mice. In addition, chronic but not acute treatment with the PDE5 inhibitor, sildenafil, synergizes with exercise training to improve performance with incremental exercise in DIO mice. These improvements with training are accompanied with increased muscle capillary flow velocity and capillary density.

The microvasculature is an important regulator of sustained metabolic activity in skeletal muscle as it controls the delivery of nutrients and other molecules that are otherwise limiting in the muscle. Endothelial function emerges from the complex interaction of hemodynamic forces, cell-cell interactions, and numerous biochemical signals. One signaling molecule, NO, catalyzed in the endothelium by eNOS, regulates vasoactivity and microvascular permeability. In healthy conditions, NO promotes vasodilation of arterioles and regulates endothelial barrier function.

Increases in vascular shear forces with physical activity enhances NO elaboration, bioavailability and vasodilation of arterioles, increasing blood flow in working muscle (10). Production of EC NO is also involved in stimulating angiogenesis (25). We tested the hypothesis that partial loss of EC eNOS-derived NO via eNOS gene heterozygosity reduces exercise training-induced improvements in microcirculatory function. We find that exercise tolerance is reduced in EC eNOS+/-mice that remain sedentary. This is consistent with the finding of decreased capillary density in EC eNOS+/-mice and suggests that eNOS is required for maintaining capillaries in the sedentary state. The training-induced increase in aerobic capacity is not blunted in eNOS+/-mice. Microvascular hemodynamics are also similar between WT and eNOS+/-mice that underwent exercise training. There are possible explanations for sustained microvascular adaptations to exercise training in eNOS+/-mice. During exercise, extracellular concentrations of lactate, K+ ions, and adenosine can increase in muscle. These molecules can cause vasorelaxation of smooth muscle that results in increased blood flow. The adaptive response to repeated exercise bouts resulting from vasoactive molecules other than NO could stimulate the production of new capillaries resulting in increased surface area for exchange and/or increase microvascular permeability to facilitate molecular exchange of nutrients during exercise. This is consistent with previous studies showing that the NOS inhibitor L-NAME does not impede exercise capacity in rodents (26). A previous study showed that whole-body eNOS+/-mice have similar submaximal exercise capacity than WT controls, despite reduced capacity to generate cGMP needed for smooth muscle vasorelaxation (27). In the present study, the EC-eNOS+/-mice are otherwise healthy. This is significant because eNOS is more likely expressed in its dimer form, which produces NO, than in the monomer form that produces superoxide (28, 29). The monomer form of eNOS is increased in disease states, which increases oxidative stress (30). Interestingly, prior findings show that mice with deletion of both eNOS alleles paradoxically have increased NO, but despite this, fail to maintain glucose levels during exercise (31). In the latter work, eNOS ablation caused a reduction in oxidative metabolism in skeletal muscle, in part, due to decreased abundance of mitochondrial complexes. Taken together, these findings suggest that in mice loss of a single eNOS allele in ECs does not impede microvascular adaptations to exercise training.

We found that increasing cGMP concentrations via chronic sildenafil administration enhances microvascular training adaptations and exercise tolerance in DIO, but not lean mice. Physiological limitations associated with vascular delivery and extraction of substances may not manifest themselves under normal circumstances but become apparent under conditions of stress. For example, healthy human subjects treated with sildenafil show no added benefits to exercise training in normoxic conditions (32). However, in conditions of hypoxia (high altitudes or experimental hypoxia), sildenafil prevents reductions in V_□_O_2max_ in healthy subjects (33-35). Excess adiposity and inflammation characteristic of DIO can be a stressor to the vasculature and to physical function. Indeed, compare to the lean state, leg blood flow and exercise tolerance are reduced in obesity. We show that chronic, but not acute, sildenafil treatment enhances aerobic work capacity and microvascular hemodynamics with training. This is similar to the effects of sildenafil on glucose uptake and insulin action. Indeed, only chronic administration of sildenafil in DIO mice increases muscle glucose uptake during an insulin clamp (1). Acute inhibition of PDE-5 increases cGMP concentrations and enhances capillary flow velocity but this does not lead to increases in exercise tolerance. One reason for the lack of improvement in aerobic work capacity during acute drug exposure is that the vasodilatory effects of sildenafil are not restricted to contracting muscle. This would predictably decrease the fraction of cardiac output in the muscle tissue because of the non-selective actions of sildenafil on vascular beds not directly involved in work production. Chronic sildenafil treatment increases proliferation of skeletal muscle capillaries. This suggests that vascular remodeling and increased density of capillaries is an important adaptation needed to reap the therapeutic benefits of PDE inhibition on skeletal muscle metabolism. Preclinical studies suggest chronic elevation in tissue cGMP levels could be beneficial in preventing or reducing fibrosis in rat models of pulmonary hypertension and streptozotocin-induced diabetes. Together these findings support that the adaptive response to elevated cGMP is an important component of its physiological efficacy with training.

### Translational implications

Clinical indications for PDE-5a inhibitors include pulmonary hypertension and erectile dysfunction. These pathologies are associated with high vascular resistance. PDE-5a inhibitors seem most effective in conditions characterized by limitations in oxygen diffusion and/or vascular delivery. This includes pathologies associated with high vascular resistance including PAD, pulmonary hypertension, and diabetes. The pathology of PAD is most sensitively manifest during physiological states characterized by repeated ischemic events. Thus, PDE-5a inhibitors combined with physical exercise are a potential mechanism for improving ambulation in patients or exercise in patients with circulatory limitations. In this preclinical study, we show that PDE-5a inhibition enhances exercise work capacity in DIO mice that are characterized with microvascular dysfunction.

In conclusion, we report that increasing cGMP with sildenafil enhances microcirculatory function and exercise work tolerance that results from training, whereas EC eNOS KD does not prevent the microcirculatory or improvements in exercise tolerance with training.

## Acknowledgements

We acknowledge the following Vanderbilt University (VU) and Vanderbilt University Medical Center (VUMC) core facilities: VU Metabolic Mouse Phenotyping Center [VMMPC (NIH DK135073; www.vmmpc.org)] and VU Cell Imaging Shared Resource. We thank Alicia Kellarakos and Carlo Malabanan for their assistance with surgical procedures.

This work is dedicated to our dear colleague Dr. David H. Wasserman.

## References

1. Ayala JE, Bracy DP, Julien BM, Rottman JN, Fueger PT, and Wasserman DH. Chronic treatment with sildenafil improves energy balance and insulin action in high fat-fed conscious mice. Diabetes. 2007;56(4):1025–33.

2. Wasserman DH. Four grams of glucose. Am J Physiol Endocrinol Metab. 2009;296(1):E11–21.

3. Carvajal JA, Germain AM, Huidobro-Toro JP, and Weiner CP. Molecular mechanism of cGMP-mediated smooth muscle relaxation. Journal of cellular physiology. 2000;184(3):409–20.

4. Fulton D, Gratton J-P, McCabe TJ, Fontana J, Fujio Y, Walsh K, et al. Regulation of endothelium-derived nitric oxide production by the protein kinase Akt. Nature. 1999;399(6736):597–601.

5. Galiè N, Ghofrani HA, Torbicki A, Barst RJ, Rubin LJ, Badesch D, et al. Sildenafil citrate therapy for pulmonary arterial hypertension. N Engl J Med. 2005;353(20):2148–57.

6. Fang J, Zhang P, Zhou Y, Chiang CW, Tan J, Hou Y, et al. Endophenotype-based in silico network medicine discovery combined with insurance record data mining identifies sildenafil as a candidate drug for Alzheimer’s disease. Nature aging. 2021;1(12):1175–88.

7. Hainsworth AH, Arancio O, Elahi FM, Isaacs JD, and Cheng F. PDE5 inhibitor drugs for use in dementia. Alzheimer’s & dementia (New York, N Y). 2023;9(3):e12412.

8. Behr-Roussel D, Oudot A, Caisey S, Coz OL, Gorny D, Bernabé J, et al. Daily treatment with sildenafil reverses endothelial dysfunction and oxidative stress in an animal model of insulin resistance. European urology. 2008;53(6):1272–80.

9. Ramirez CE, Nian H, Yu C, Gamboa JL, Luther JM, Brown NJ, et al. Treatment with Sildenafil Improves Insulin Sensitivity in Prediabetes: A Randomized, Controlled Trial. The Journal of clinical endocrinology and metabolism. 2015;100(12):4533–40.

10. Sarelius I, and Pohl U. Control of muscle blood flow during exercise: local factors and integrative mechanisms. Acta Physiol (Oxf). 2010;199(4):349–65.

11. Hamburg NM, and Balady GJ. Exercise rehabilitation in peripheral artery disease: functional impact and mechanisms of benefits. Circulation. 2011;123(1):87–97.

12. Xia N, Horke S, Habermeier A, Closs EI, Reifenberg G, Gericke A, et al. Uncoupling of Endothelial Nitric Oxide Synthase in Perivascular Adipose Tissue of Diet-Induced Obese Mice. Arterioscler Thromb Vasc Biol. 2016;36(1):78–85.

13. Gamez-Mendez AM, Vargas-Robles H, Rios A, and Escalante B. Oxidative Stress-Dependent Coronary Endothelial Dysfunction in Obese Mice. PLoS One. 2015;10(9):e0138609.

14. Ketonen J, Pilvi T, and Mervaala E. Caloric restriction reverses high-fat diet-induced endothelial dysfunction and vascular superoxide production in C57Bl/6 mice. Heart and vessels. 2010;25(3):254–62.

15. Elefanty AG, Begley CG, Hartley L, Papaevangeliou B, and Robb L. SCL expression in the mouse embryo detected with a targeted lacZ reporter gene demonstrates its localization to hematopoietic, vascular, and neural tissues. Blood. 1999;94(11):3754–63.

16. Sinclair AM, Gottgens B, Barton LM, Stanley ML, Pardanaud L, Klaine M, et al. Distinct 5’ SCL enhancers direct transcription to developing brain, spinal cord, and endothelium: neural expression is mediated by GATA factor binding sites. Dev Biol. 1999;209(1):128–42.

17. Garrido-Martin EM, Nguyen HL, Cunningham TA, Choe SW, Jiang Z, Arthur HM, et al. Common and distinctive pathogenetic features of arteriovenous malformations in hereditary hemorrhagic telangiectasia 1 and hereditary hemorrhagic telangiectasia 2 animal models--brief report. Arterioscler Thromb Vasc Biol. 2014;34(10):2232–6.

18. Gothert JR, Gustin SE, van Eekelen JA, Schmidt U, Hall MA, Jane SM, et al. Genetically tagging endothelial cells in vivo: bone marrow-derived cells do not contribute to tumor endothelium. Blood. 2004;104(6):1769–77.

19. Williams IM, Valenzuela FA, Kahl SD, Ramkrishna D, Mezo AR, Young JD, et al. Insulin exits skeletal muscle capillaries by fluid-phase transport. J Clin Invest. 2018;128(2):699–714.

20. Williams IM, McClatchey PM, Bracy DP, Bonner JS, Valenzuela FA, and Wasserman DH. Transendothelial Insulin Transport is Impaired in Skeletal Muscle Capillaries of Obese Male Mice. Obesity (Silver Spring, Md). 2020;28(2):303–14.

21. McClatchey PM, Williams IM, Xu Z, Mignemi NA, Hughey CC, McGuinness OP, et al. Perfusion controls muscle glucose uptake by altering the rate of glucose dispersion in vivo. Am J Physiol Endocrinol Metab. 2019;317(6):E1022–E36.

22. McClatchey PM, Mignemi NA, Xu Z, Williams IM, Reusch JEB, McGuinness OP, et al. Automated quantification of microvascular perfusion. Microcirculation. 2018;25(6):e12482.

23. Wasserman DH, Wang TJ, and Brown NJ. The Vasculature in Prediabetes. Circulation research. 2018;122(8):1135–50.

24. Padilla J, Manrique-Acevedo C, and Martinez-Lemus LA. New insights into mechanisms of endothelial insulin resistance in type 2 diabetes. Am J Physiol Heart Circ Physiol. 2022;323(6):H1231–H8.

25. Matsunaga T, Weihrauch DW, Moniz MC, Tessmer J, Warltier DC, and Chilian WM. Angiostatin inhibits coronary angiogenesis during impaired production of nitric oxide. Circulation. 2002;105(18):2185–91.

26. Wojewoda M, Przyborowski K, Sitek B, Zakrzewska A, Mateuszuk L, Zoladz JA, et al. Effects of chronic nitric oxide synthase inhibition on V’O(2max) and exercise capacity in mice. Naunyn Schmiedebergs Arch Pharmacol. 2017;390(3):235–44.

27. Kojda G, Cheng YC, Burchfield J, and Harrison DG. Dysfunctional regulation of endothelial nitric oxide synthase (eNOS) expression in response to exercise in mice lacking one eNOS gene. Circulation. 2001;103(23):2839–44.

28. Andrew PJ, and Mayer B. Enzymatic function of nitric oxide synthases. Cardiovascular research. 1999;43(3):521–31.

29. Xu H, Shi Y, Wang J, Jones D, Weilrauch D, Ying R, et al. A heat shock protein 90 binding domain in endothelial nitric-oxide synthase influences enzyme function. The Journal of biological chemistry. 2007;282(52):37567–74.

30. Schulz E, Jansen T, Wenzel P, Daiber A, and Münzel T. Nitric oxide, tetrahydrobiopterin, oxidative stress, and endothelial dysfunction in hypertension. Antioxid Redox Signal. 2008;10(6):1115–26.

31. Lee-Young RS, Ayala JE, Hunley CF, James FD, Bracy DP, Kang L, et al. Endothelial nitric oxide synthase is central to skeletal muscle metabolic regulation and enzymatic signaling during exercise in vivo. Am J Physiol Regul Integr Comp Physiol. 2010;298(5):R1399–408.

32. Kressler J, Stoutenberg M, Roos BA, Friedlander AL, Perry AC, Signorile JF, et al. Sildenafil does not improve steady state cardiovascular hemodynamics, peak power, or 15-km time trial cycling performance at simulated moderate or high altitudes in men and women. Eur J Appl Physiol. 2011;111(12):3031–40.

33. Faoro V, Lamotte M, Deboeck G, Pavelescu A, Huez S, Guenard H, et al. Effects of sildenafil on exercise capacity in hypoxic normal subjects. High altitude medicine & biology. 2007;8(2):155–63.

34. Richalet JP, Gratadour P, Robach P, Pham I, Dechaux M, Joncquiert-Latarjet A, et al. Sildenafil inhibits altitude-induced hypoxemia and pulmonary hypertension. Am J Respir Crit Care Med. 2005;171(3):275–81.

35. Ghofrani HA, Reichenberger F, Kohstall MG, Mrosek EH, Seeger T, Olschewski H, et al. Sildenafil increased exercise capacity during hypoxia at low altitudes and at Mount Everest base camp: a randomized, double-blind, placebo-controlled crossover trial. Annals of internal medicine. 2004;141(3):169–77.

